# *OsPSTOL* but not *TaPSTOL* can play a role in nutrient use efficiency and works through conserved pathways in both wheat and rice

**DOI:** 10.1101/2022.11.02.514825

**Authors:** Matthew Milner, Sarah Bowden, Melanie Craze, Emma J. Wallington

## Abstract

There is a large demand to reduce inputs for current crop production, particularly phosphate and nitrogen inputs which are the two most frequently added supplements to agricultural production. Gene characterization is often limited to the native species from which it was identified, but may offer benefits to other species. To understand if *OsPSTOL*, a gene identified from rice which improves tolerance to low P growth conditions, might improve performance and provide the same benefit in wheat, *OsPSTOL* was transformed into wheat and expressed from a constitutive promoter. The ability of *OsPSTOL* to improve nutrient acquisition under low phosphate or low nitrogen was evaluated. Here we show that *OsPSTOL* works through a conserved pathway in wheat and rice to improve yields under both low phosphate and low nitrogen. This increase is yield is mainly driven by improved uptake from the soil driving increased biomass and ultimately increased seed number, but does not change the concentration of N in the straw or grain. Overexpression of *OsPSTOL* in wheat modifies N regulated genes to aid in this uptake whereas the putative homolog *TaPSTOL* does not suggesting that expression of *OsPSTOL* in wheat can help to improve yields under low input agriculture.

## Introduction

There is a strong desire to reduce inputs in agriculture to make large scale farming more sustainable and lessen our impact on the surrounding environment. Phosphorous and nitrogen are added in large quantities as chemical fertilizers in modern agricultural practices. Both elements are essential macronutrients which are required for all major developmental processes in plants and are key for both completing the life cycle and maintaining food production. Their mobility within the soil, mechanisms of uptake and the plant scavenge response to their limitation differ (Marschner, 1995).

Recent advances in our understanding of the molecular mechanisms by which different plant species adapt to different abiotic stresses, including the regulation and expression of key genes and growth habits have enabled the design of more effective breeding strategies to produce highly resilient crops (Gamuyao *et al*., 2012; Magalhaes *et al*., 2007; Milner *et al*., 2022; Xu *et al*., 2006). A number of the underlying genes involved in the response of plants to low P and N are highly conserved and play similar roles in a number of diverse plant species including both model system and crop species (Borah *et al*., 2018; Milner *et al*., 2022). Thus, a greater understanding of the pathways involved, the key genes involved in P and N acquisition and signaling will allow breeders and plant molecular biologists to develop more efficient crops. Identification of these conserved pathways and genes from “model” organisms, and the subsequent transfer of this knowledge to crop species, would therefore allow farmers to optimize fertilizer use worldwide, resulting in increased efficiency of food production with lower environmental cost.

One such gene, believed to be important in increased nutrient use efficiency, is the recent identification of a key gene under the PUP1 locus from rice. The gene has been shown to be a serine/threonine receptor-like kinase, named Phosphate Starvation Tolerance 1 (PSTOL), is a major gene involved in tolerance to growth on low P soils (Chin *et al*., 2011; Gamuyao *et al*., 2012; Heuer *et al*., 2009; Vigueira *et al*., 2016). The PUP1 locus was originally identified in an upland variety of rice, Kasalath, yet is absent from most rice cultivars including the current reference japonica variety Nipponbare and indica variety 93-11 (Vigueira *et al*., 2016; Wissuwa *et al*., 2002). Rice varieties which have this genomic introgression containing a functional PSTOL gene, show increased biomass, increased root growth, increased tiller number and yield increases of up to 30% when grown under low P conditions whereas no deleterious consequences were seen when grown under normal soil fertility conditions (Chin *et al*., 2011; Gamuyao *et al*., 2012). The identification of OsPSTOL and its role in helping rice tolerate low P conditions has led to the belief that one could increase the PUE by incorporation of PSTOL type genes into many crop species. However when OsPSTOL was first identified the authors also suggested that other elements may also benefit from in increased root growth and make the plant more efficient to low nitrogen (N) conditions and drought (Gamuyao *et al*., 2012), but as yet no data been presented to substantiate the possibility of a role outside of PUE.

Others have identified homologous PSTOL-like genes in both maize and sorghum, based upon QTL analysis and sequence homology (Azevedo *et al*., 2015; Bernardino *et al*., 2019; Hufnagel *et al*., 2014). Further evidence of the role of PSTOL-like genes in PUE has been supported through QTL mapping rather than direct molecular characterization of candidate genes. Criteria to identify other potential

PSTOL like genes has included identification of protein domains such as ATP kinase domains and on DNA sequence conservation meeting certain bioinformatic cut-offs for genes underlying these QTL. Some of these homologues appear to have differences in their gene structure such as number and length of introns and UTRs. Despite the lack of a highly conserved gene in most plant species, it is critical to understand whether other *PSTOL* genes exist in other important crop species and if they can be exploited. Our previous study identified a putative wheat PSTOL gene *(TaPSTOL)* and characterized its role in PUE and other important agronomic phenotypes. As the overexpression of TaPSTOL was not able to replicate the yield increase seen with overexpression of *OsPSTOL* in rice, we suggested that either wheat lacked the other genes downstream of PSTOL to increase yields or a critical domain within the coding sequence itself was the reason for the lack of increased yields.

Here, we set out to test whether the differences seen in a plants ability to maintain yield on low P is due to the presence of the *OsPSTOL* or *TaPSTOL* genes *per se* or if other parts of an unknown pathway underlie the differences seen in yield increases. We also test whether expression of *OsPSTOL* can overcome other nutrient deficiencies to see if the presence of *OsPSTOL* is only beneficial to low phosphate conditions.

## Methods

### Cloning

The *OsPSTOL* coding sequence from EMBL (AB458444.1) was synthesized with a ribosomal binding site, CCACC, immediately upstream of the ATG start codon and flanked with gateway attL1/attL2 sites (Genewiz). The synthesized sequence was then recombined into the binary vector pSc4ActR1R2 using a Gateway LR Clonase II kit (Thermofisher) to create the plasmid pMM007 for stable wheat transformation. In pMM007 *OsPSTOL* is driven by the OsActin promoter and terminated by the *A. tumefaciens* nopaline synthase terminator (tNOS).

### Wheat Transformation

Wheat cv. Fielder plants were grown in controlled environment chambers (Conviron) at 20 °C day/15 °C night with a 16 h day photoperiod (approximately 400 μE m^-⍰2^ s^-⍰1^). Immature seeds were harvested for transformation experiments at 14–20 days post-anthesis. Isolated immature wheat embryos were co-cultivated with *Agrobacterium tumefaciens* for 2 days in the dark [35]. Subsequent removal of the embryonic axis and tissue culture was performed as previously described [36]. Individual plantlets were hardened off following transfer to Jiffy-7 pellets (LBS Horticulture), potted up into 9 cm plant pots containing M2 compost plus 5 g/l slow-release fertilizer (Osmocote Exact 15:9:9) and grown on to maturity and seed harvest in controlled environment chambers. *TaPSTOL* overexpression line OE-1 was also grown for comparison (Milner *et al*., 2018).

### DNA analysis of transformed wheat plants

Wheat plantlets which regenerated under G418 selection in tissue culture were transferred to Jiffy-7 pellets and validated using an nptII copy number assay relative to a single copy wheat gene amplicon, GAMyb for wheat, normalized to a known single copy wheat (Milner *et al*., 2018). Primers and Taqman probes were used at a concentration of 10 μM in a 10 μl multiplexed reaction using ABsolute Blue qPCR ROX mix (Thermofisher) with the standard run conditions for the ABI 7900 HT for wheat. The relative quantification, ΔΔ^ct^, values were calculated to determine nptII copy number in the T_0_ and subsequent generations. Primers for the copy number determination are listed in Suppl. Table 1. Homozygous and null transgenic lines were identified on the basis of copy number for the selection gene and segregation analysis in the T_1_ generation. Null segregates were used for further study and referred to as WT Fielder.

### Plant growth conditions

Plants were grown in a controlled growth chamber under 16 h light with 20 °C/15 °C day/night temperatures for all conditions tested. To study low P conditions WT and transgenic wheat lines were grown under in sand and fertilized twice a week with nutrient solution (Magnavaca *et al*., 1987) containing 3 μM KH_2_PO_4_, 1.3 mM NH_4_NO_3_, 3.52 mM Ca(NO_3_)_2_, 0.58 mM, KCl, 0.58 mM K_2_SO_4_, 0.56 mM KNO_3_, 0.86 mM Mg(NO_3_)_2_ 0.13 mM H_3_BO_3_, 5 μM MnCl_2_, 0.4 μM Na_2_MoO_4_, 10 μM ZnSO_4_, 0.3 μM CuSO_4_, Fe(NO_3_)_3_ and 2 mM MES (pH 5.5) twice a week until maturity. For low N conditions wheat and plants were grown on TS5 low fertility soil to control total nitrogen with a starting nitrogen level of 0.1 mg/l (Bourne Amenity, Kent, UK). Ammonium nitrate was then added to reach a final concentration in the pots equivalent to field fertilizer application of 70, 140 or 210 kg N /ha which equates to 23.3, 46.6, or 70⍰mg/ 1 L pot. Each pot also received 4.2⍰mg Ca, 2.7⍰mgl3K, 0.62⍰mg Mg, 0.04⍰mg⍰P, 0.56⍰mg⍰S, 0.008⍰mg B, 0.13⍰mg Fe, 0.015⍰mg Mn, 0.0012⍰mg Cu, 0.0024 Mo, 0.0045 Zn, 0.00⍰mg Na, and 0.63⍰mg Cl per pot added as Magnavaca solution which was added 10 days after sowing.

### Whole plant measurements

Total dry shoot weight, seed weight (yield per plant), seed number, seed size, tiller number. Biological replicates each contained at least 14 plants per line and were grown until seed maturation. Tissues were allowed to dry for a further two weeks before harvesting.

### N tissue measurements

Samples were measured using the Dumas method. The samples were dried for 17 hours at 100°C and then milled on a 1mm hammer mill. Prior to testing the sample were dried at 104°C for 3 hours and 1g of sample was loaded on the instrument (Leco TruMacN Dumas gas analyser), following the manufacturer’s instructions. Samples were converted to gases by heating in a combustion tube at 1150°C. Interfering components are removed from the resulting gas mixture. The nitrogen compounds in the gas mixture or a representative part of the mixture, are converted to molecular nitrogen which is quantitatively determined by a thermal conductivity detector. The nitrogen content is then calculated by a microprocessor.

### RNA expression analysis

Wheat seedlings were grown for seven days in 2.2 L pots containing Magnavaca solution as listed above and amended with 3 μM KH_2_PO_4_ for low P treatments or omitting 1.3 mM NH_4_NO_3_, and 3.52 mM Ca(NO_3_)_2_ for low N treatment Plants were grown for an additional 7 days before harvesting tissue and separating the samples into root and shoot tissues for analysis. Total RNA was isolated from both roots and shoots for each treatment using a RNeasy Kit (Qiagen) and treated with DNasel (Thermofisher) prior to cDNA synthesis from 500 ng of total RNA using Omniscript RT Kit (Qiagen). The cDNA was diluted 1:2 with water and 0.5 μL was used as template in each RT-PCR reaction. Expression levels were quantified by quantitative PCR in triplicate reactions from three biological replications using SYBR Green JumpStartTaq ReadyMix (SIGMA) with the standard run conditions for the ABI 7900 HT. OsPSTOL expression was compared to TaUbi. Primers used for amplification of transcripts are listed in Suppl. Table 1.

### ^15^N uptake

To measure N uptake a similar protocol as previously reported (Milner *et al*., 2022), briefly roots from 2 week old seedlings were exposed to ^15^NH_4_^15^NO_3_ for 10 min, then washed in 0.1 mM CaSO_4_ for 2 min, harvested separated into root and shoot tissue, and dried at 70°C for 48 h before grinding. Dried tissue was then placed in 2mL microfuge tubes with 2 × 5mm diameter stainless steel beads and shaken in a genogrinder until a fine powder was obtained. Dried and ground samples were carefully weighed (0.5 mg) into tin capsules, sealed and loaded into the auto-sampler. Samples were analyzed for percentage carbon, percentage nitrogen, ^12^C/^13^C (δ^13^C) and ^14^N/^15^N (δ^15^N) using a Costech Elemental Analyzer attached to a Thermo DELTA V mass spectrometer in continuous flow mode. The excess ^15^N was calculated based on measurements of δ^15^N and tissue N%. The absolute isotope ratio (R) was calculated for labelled samples and controls, using R_standard_ (the absolute value of the natural abundance of ^15^N in atmospheric N_2_; 1).

1. (1) R_sample or control_ = [(δ^15^N/1000)+1] x R_standard_ Then, molar factional abundance (F) and mass-based factional abundance (MF) were calculated (2,3,4)
2. F= R_sample or control_/(R_sample or control_+1)
3. MF= (F x 15) x /[(F x 15)+ ((1-F)x 14)]
4. ΔMF = MF_sample_ - MF_control_ The excess ^15^N in mg in a total tissue was calculated as in (5)
5. Excess ^15^N (g)= ΔMF x Tissue DW (g) x Tissue N%/100

### Chlorophyll measurements

Leaf chlorophyll content was determined using the method developed by Hiscox et al. (Hiscox and Israelstam, 1979). Chlorophyll was extracted from 100mg of fresh leaf tissue from six plants into 20mL DMSO, mixed for on a rotary shaker for 30 mins and then placed at 4°C overnight. Chlorophyll measurements were taken at 645 and 663 nm (spectrophotometer Jenway model 7315, Staffordshire, UK).

### Carbon assimilation measurements

An LI-6800 portable photosynthesis infrared gas analyzer system (LI-COR) equipped with a multiphase flash fluorimeter was used to assess physiological differences for photosynthetic parameters between transgenic and WT wheat plants. Measurements were taken on the fourth leaf of plants grown on TS5 low fertility soil (Bourne Amenity, Kent, UK). Ammonium nitrate or K_2_PO_4_ was then added to reach a final concentration in the pots equivalent to field fertilizer application of 70 kg N /ha for low N or 50 kg P /ha for Phosphate deficiency. Plants were grown in a climate-controlled chamber with supplemented light (250 μmol.m^-2^/s^-1^) for a 16hr day and 20°C/15°C day night temperatures for wheat. All measurements were also normalized for the amount of area of the measuring disk. Measurements were carried out on consecutive days between 1 and 8 h post dawn, measuring three plants total selected at random from each treatment per day.

## Results

### Growth on low P

To understand if the lack of a response of plants expressing *TaPSTOL* was due to some other mitigating factor other than OsPSTOL wheat plants were transformed with *OsPSTOL* driven by the OsActin promoter as described previously for *TaPSTOL* (Milner *et al*., 2018). Three highly expressing *OsPSTOL* wheat transgenic lines were compared to the previously created *TaPSTOL* overexpressing line (OE-1), and WT Fielder for their ability to improve growth under low P conditions (Milner *et al*., 2018). As seen in figure 1, the three transgenic wheat lines expressing *OsPSTOL* showed increased biomass and increased yield compared to either TaOE-1 or WT wheat plants when grown under low P conditions (p val 0). When compared to a null segregant, lines expressing *OsPSTOL* but not *TaPSTOL* showed increased biomass and yields when grown on low P soil (Figure 1). This includes increased yields of 29 to 47% compared with the null segregant. The yield increase is most likely due to higher biomass production leading to higher seed set as the number of seeds per plant was significantly higher in wheat lines expressing *OsPSTOL* (Figure 1C). There was a significant difference found in yields between TaOE-1 and WT wheat lines in their yield on low P grown plants, but above ground biomass produced per plant was not significantly different, similar to previous reports (Milner *et al*., 2018).

**Figure 1:**
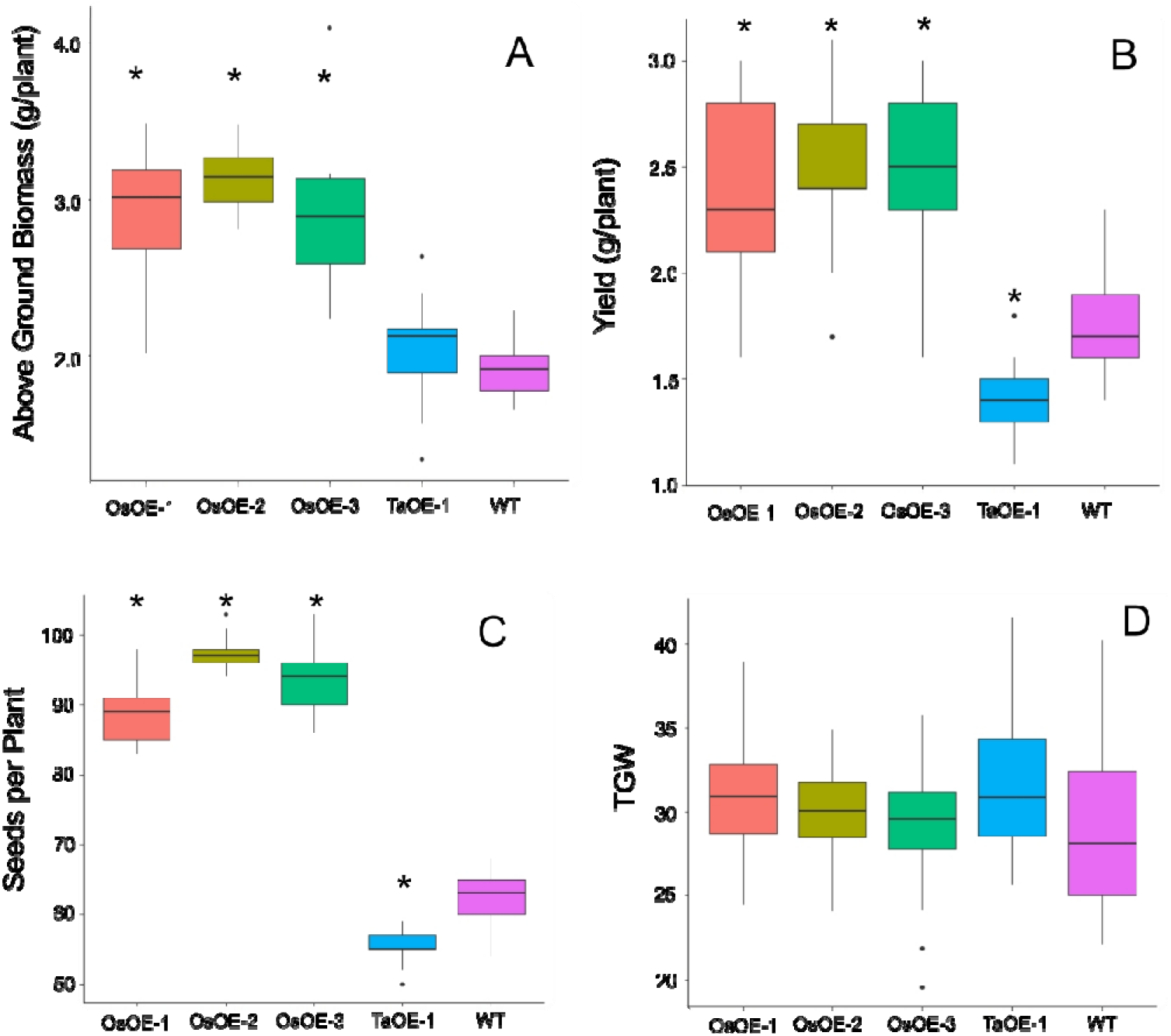
Agronomic traits of transgenic wheat plants expressing either OsPSTOL or TaPSTOL grown under low P conditions. A) Above ground biomass B) yield per plant C) seeds per plant D) thousand grain weight (TGW). A star indicates a significant differnce (p val <0.05) between an overexpression line and WT.

### Growth on low N

To determine if expression of *OsPSTOL* can lead to increased yield under other nutrient limiting conditions such a low N, the three transgenic wheat lines were grown under three N levels to study the transgenes effect on growth. The three levels of N were equivalent to 70, 140 and 210 kg/ha N levels in soil. As shown with growth in low P growth conditions, increases in both biomass and yield were seen in plants expressing *OsPSTOL*, but not *TaPSTOL* (Figure 2). This difference is mainly driven by growth under the higher N levels as no significant differences were seen in OsOE-1 or OsOE-2, either in above ground biomass or per plant yield at the lowest N level (p vals, 0.07 and 0.33). At higher N levels (140 and 210) significant differences were seen in OE-2 and OE-3 for both traits. As seen previously with growth on low P, the gains in yield were mainly driven by increased biomass leading to increased seed set and consequently higher yields.

**Figure 2:**
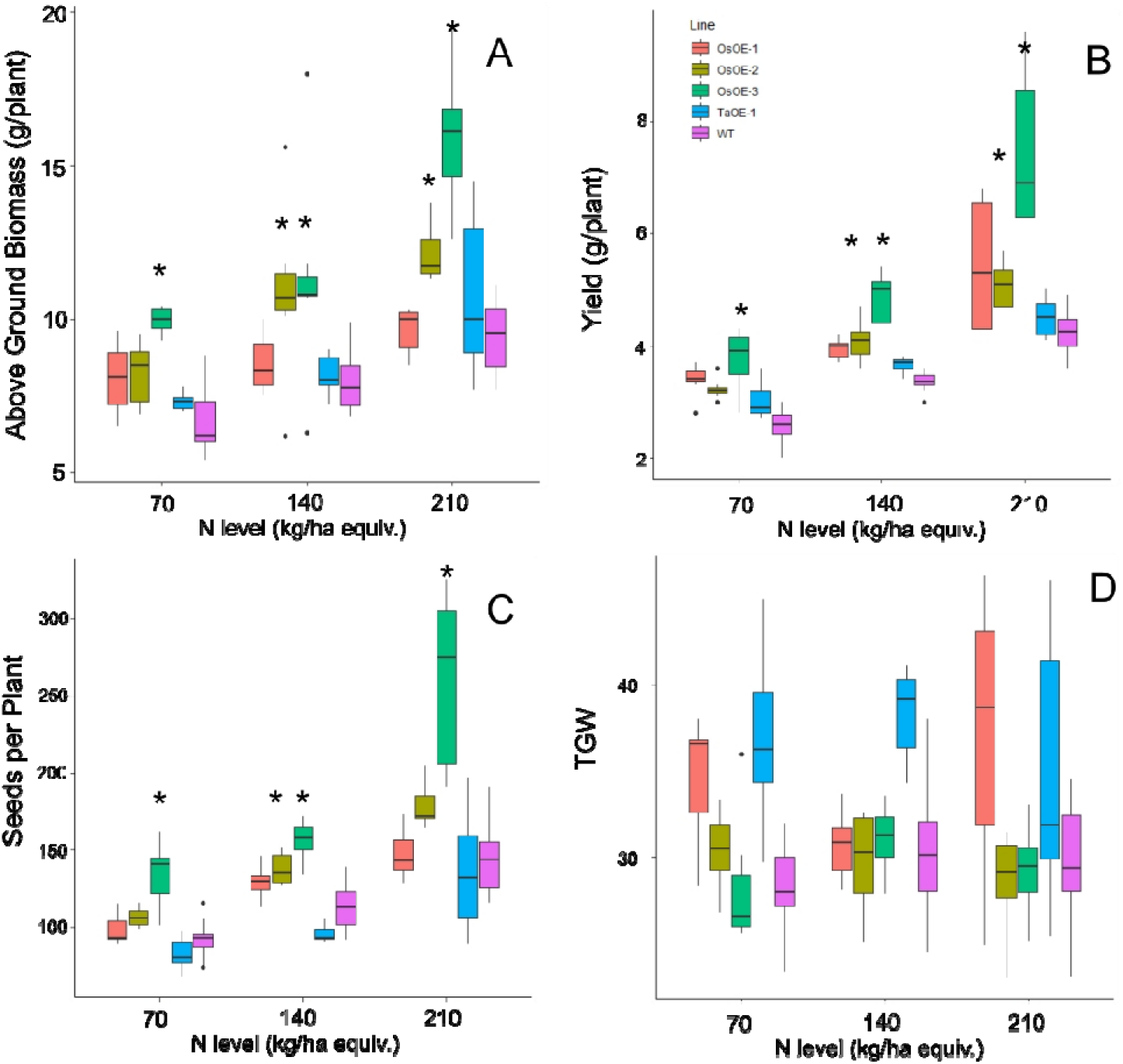
Agronomics of Os and TaPSTOL lines grown under a range of N levels. A) Above ground biomass B) Yield per plant C) seeds per plant D) Thousand Grain Weight (TGW). A star indicates a significant difference between an overexpression line and WT (pval < 0.05).

N levels in the grain and leaf tissue were measured by DUMAS to determine whether many of the same phenotypes of overexpression of *OsPSTOL* were seen under low N as when grown under low P conditions. No significant differences were seen in the straw N levels for any of the OsOE lines or TaOE line compared to WT (Figure 3). In the grain a significant difference in N content could be seen with significant differences in N concentrations for two of three OsPSTOL overexpression lines, OsOE-1 and OsOE-3, with OsOE-2 was just outside the 0.05 cutoff for significance relative to WT (p vals 0.03, 0.0005, 0.07). No significant difference was observed in N content of the TaOE-1 lines relative to WT wheat in the grain under any N level tested (p val. 0.31). Only OsOE-3 had significantly more N in the grain under 210 kg/ha treatments (p val 0.02) and no other direct comparison between a line at the same treatment was significantly different.

**Figure 3:**
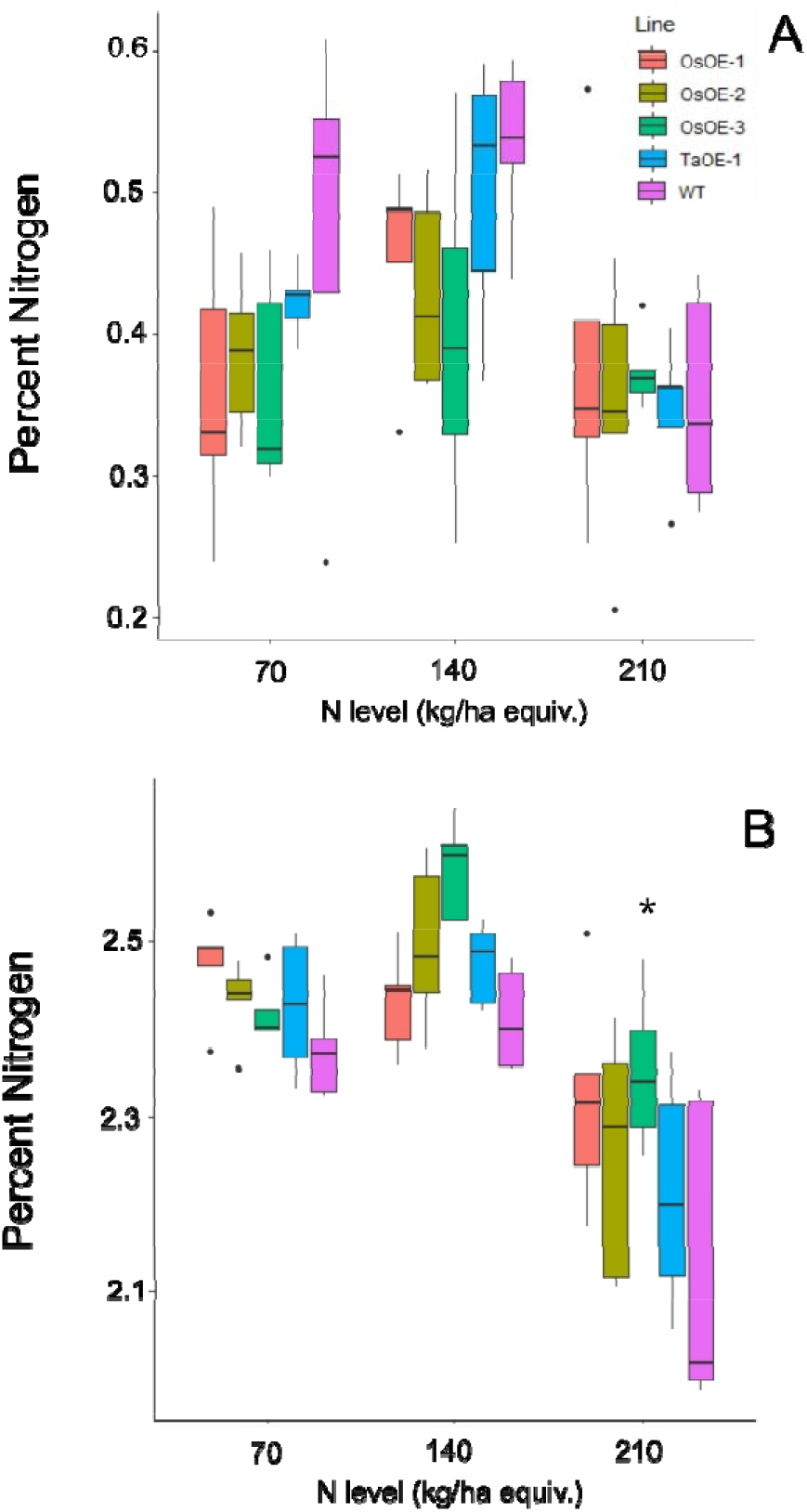
N content of Os and TaPSTOL lines grown under a range of N levels. A) Percent nitrogen in the flag leaf B) Percent nitrogen in the grain. A star indicates a significant difference between an overexpression line and WT (pval < 0.05).

To understand if the N deficiency increased transcript levels of *OsPSTOL* or if transcript levels are higher only in response to low P, we tested expression of both *OsPSTOL* in the rice variety Kasalth by qPCR and *TaPSTOL* expression in Fielder. As seen in Figure 4, *OsPSTOL* can be activated by low N growth conditions although this change in transcript level is lower than that of low P conditions (Figure 4A). But both low N and low P growth conditions significantly increased *OsPSTOL* expression in the roots of Kasalath rice plants. *TaPSTOL* expression was also measured in Roots of wheat cv. Fielder via qPCR and no change in transcript levels could be seen in response to N level relative to replete, although low P did significantly increase *TaPSTOL* expression (Figure 4B).

**Figure 4:**
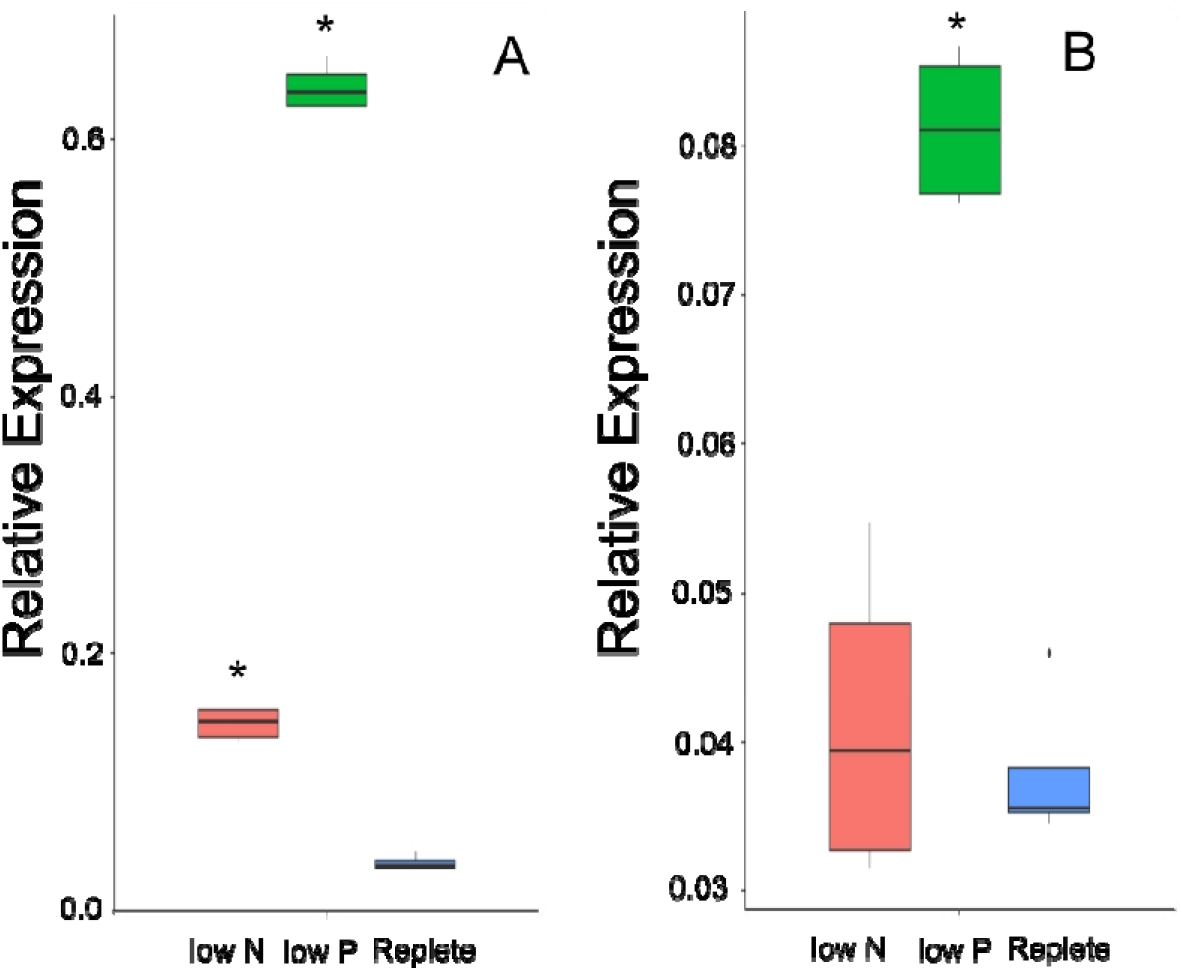
Transcript levels of OsPSTOL or TaPSTOL in the roots under different nutrient levels in rice and wheat. Expression values shown are PSTOL transcript levels relative to transcript levels of *OsUbi* for rice (A) and *TaUbi* for wheat (B) plants grown under low nitrogen (low N), low phosphate (low P) and replete nutrient levels in hydroponic solution. Data shown as the mean values (central line), lower and upper quartiles (box), minimum and maximum values (whiskers), and outliers as individual points. The statistical analysis was performed with ANOVA and post hoc Tukey test, letters correspond to significant differences between transcript levels of either line under either treatment (p⍰<⍰0.05).

### LICOR

To understand how *OsPSTOL* activates enhanced biomass we measured the rate at which plants were able to fix carbon under low P or low N conditions. When measuring C assimilation rates under low N and low P no significant differences could be seen the levels of C being fixed under the given growth conditions or replete conditions (Figure 5A). This was further supported by the levels of chlorophyll in the leaves of the treated plants as again no significant differences were seen between lines under each condition tested (Figure 5B).

**Figure 5:**
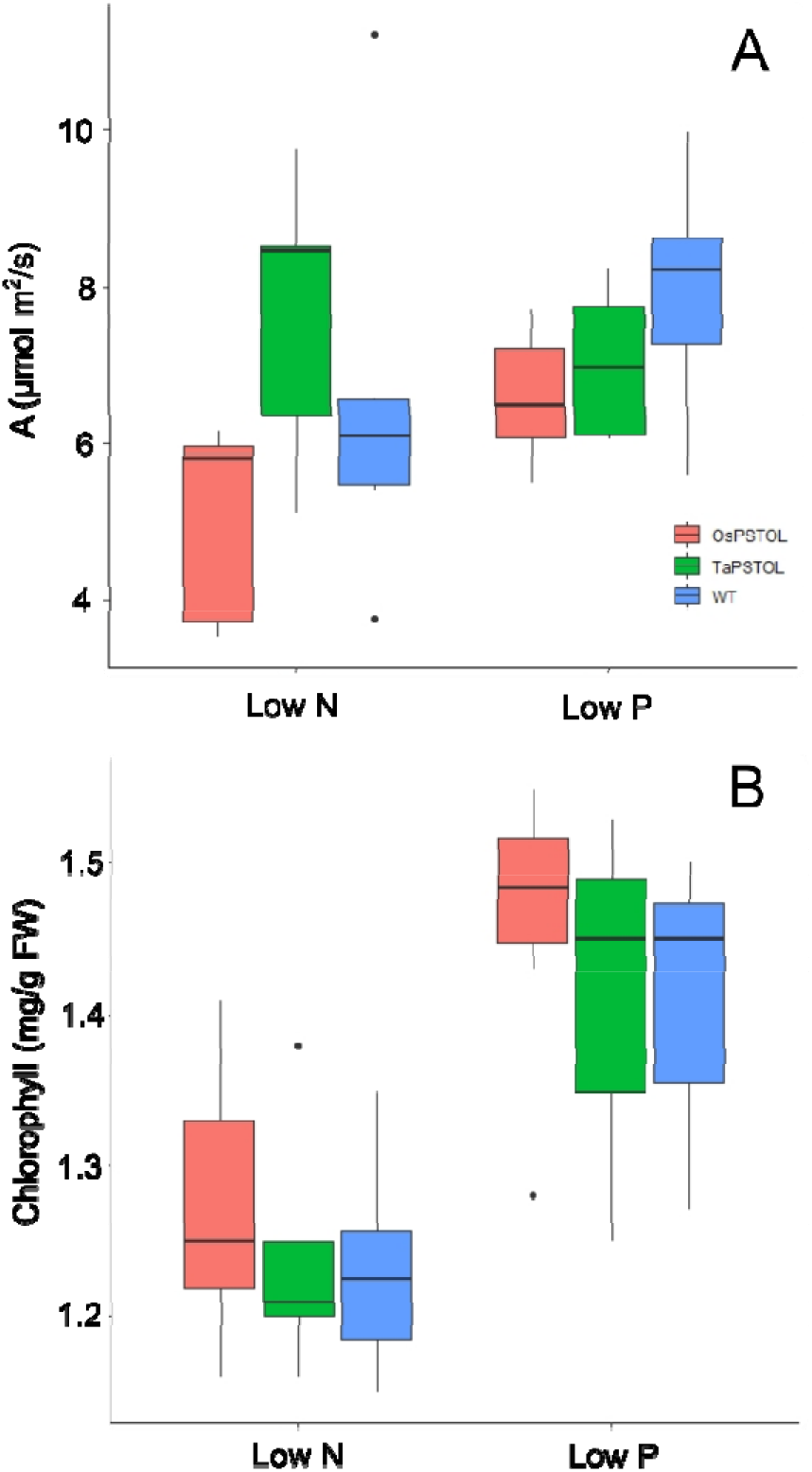
Spot measurements of C assimilation and chlorophyll content in plants overexpressing either OsPSTOL or TaPSTOLOE-1 relative to WT grown under low nitrogen (low N) or low phosphate (low P) conditions. A) Spot measurements of C assimilation grown in growth chamber conditions which include a light intensity of 250⍰μmol⍰m^2^⍰/s and CO_2_ level of 400 ppm. B) Chlorophyll content of plants grown under low nitrogen (low N) or low phosphate (low P) conditions.

### N Uptake and Expression of N related genes

To further understand how expression of *OsPSTOL* in wheat is allowing for increased growth under a range of N conditions we studied direct uptake of N in the form of ammonium nitrate, with each N atom labeled as ^15^N. When comparing rates of uptake in the roots of N deprived plants a significant difference in the amount of N being taken up per g of tissue was observed (Figure 6). This was found only in plants overexpressing *OsPSTOL* as no significant differences in uptake were seen in the roots of plants overexpressing *TaPSTOL* relative to WT roots (p val 0.13).

**Figure 6:**
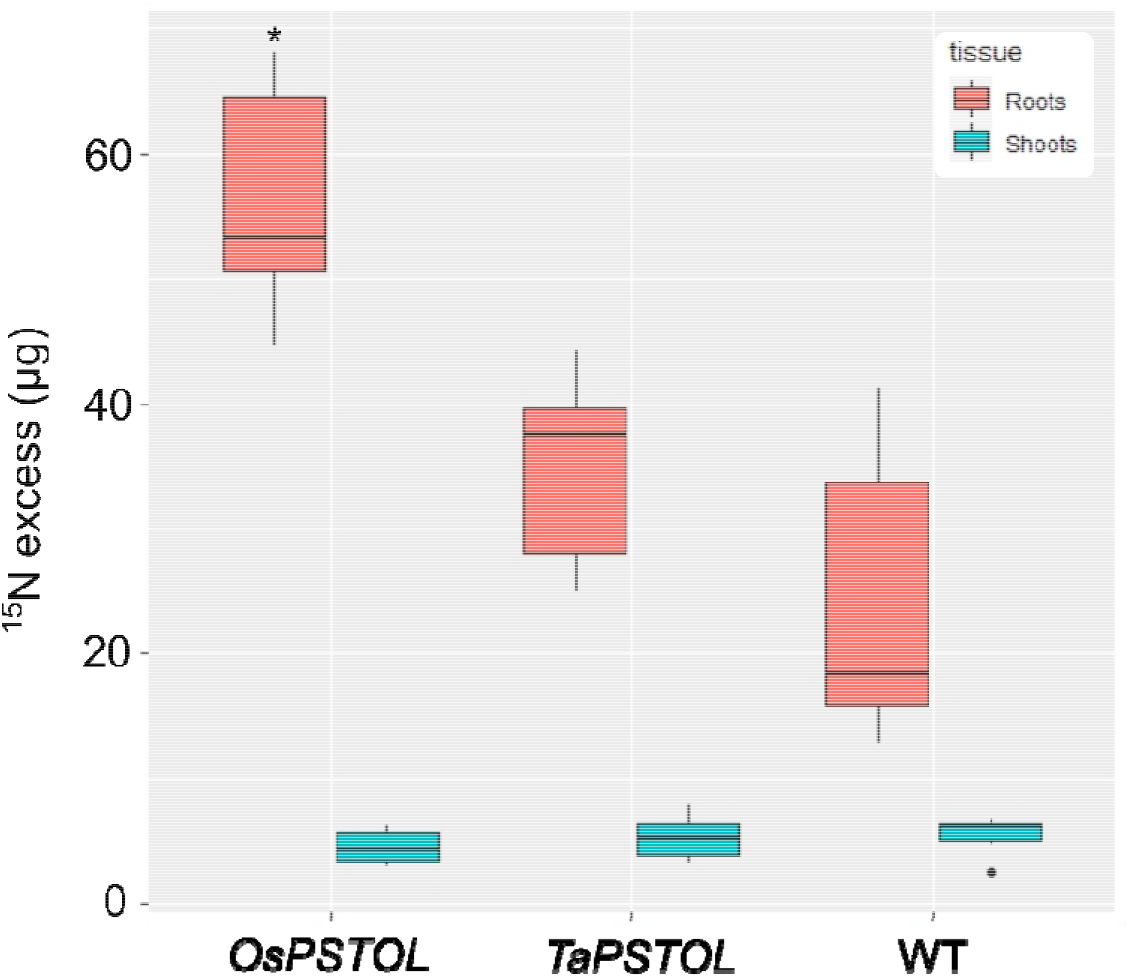
Uptake of ^15^N in the roots and shoots of overexpression lines expressing either *OsPSTOL* (OsOE-3) or *TaPSTOL* (TaOE-1) relative to WT. Data are shown as mean values (central line), lower and upper quartiles (box), minimum and maximum values (whiskers), and outliers as individual points. The statistical analysis was performed with ANOVA and post hoc Tukey test. Asterisks indicate a significant difference (p⍰<⍰0.05) between WT and an OE line in the same tissue.

The expression of four genes known to be differentially regulated under low N conditions in wheat were selected to test the N-responsiveness at the transcript level of the OsOE line (OE-3), TaOE-1 relative to WT (Figure 7) (Buchner and Hawkesford, 2014). The genes selected encode the high and low-affinity N uptake transporters *(TaNRT2.1* and *TaNRT1*), an N transporter involved in N translocation through the plant *(TaNPF7.1)*, and glutamate dehydrogenase *(TaGDH2)*, an enzyme involved in N remobilization in response to limiting carbon ©. Expression of a number of the genes were differentially expressed in the transgenic lines relative to WT Fielder. Measurements of *TaGDH2* transcript levels showed no significant differences in the roots of OsOE-3, TaOE-1 or WT. There was a significant increase in *TaGDH2* transcript levels seen in shoots of OsOE-3 compared to WT (pval <0.001). For *TaNRT1*, transcripts in roots of OsOE-3 showed significantly lower expression under replete conditions (p val < 0.001). Both OsOE3-3 and TaOE-1 showed higher transcript levels of *TaNRT1* in the shoots under both low N and replete conditions (pval <0.001). Analysis of *TaNPF7.1* transcript levels in the roots showed similar expression to that of *TaNRT1* with only OsOE-3 showing significantly lower expression than WT (p val <0.001). In the shoots *TaNPF7.1* transcript levels were higher under low N growth conditions in OsOE-3 and TaOE-1 relative to WT, but no difference in expression was seen in plants grown under replete N levels (p val <0.001). The high affinity uptake transporter *TaNRT2.1* showed significantly higher transcript levels in the roots of both OsOE-3 and TaOE-1 under both low N and replete growth conditions (p val <0.01). No significant difference in the transcript levels of *TaNRT2.1* was seen in the shoots of either overexpression line relative to WT under either N growth conditions.

**Figure 7:**
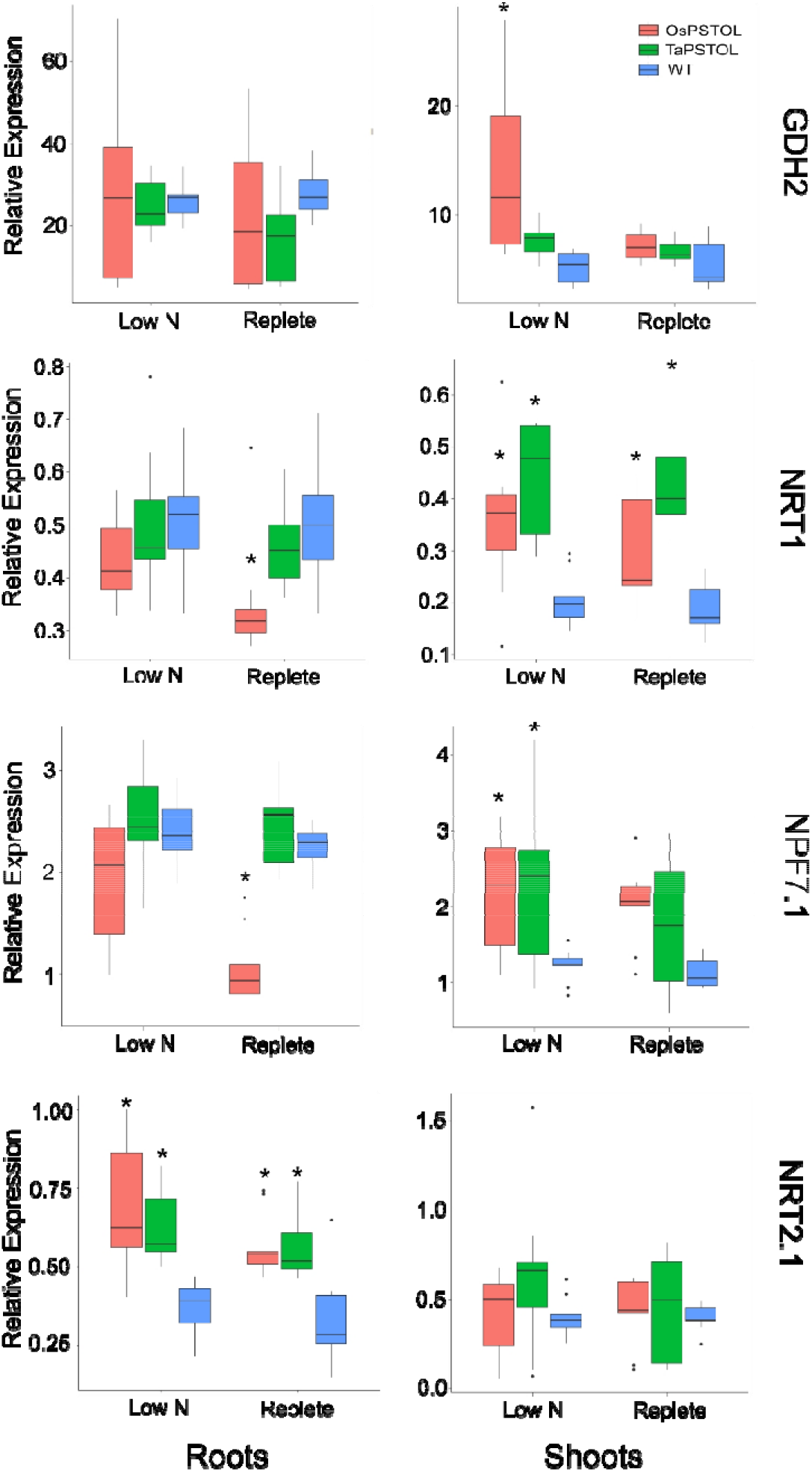
Transcript levels for N regulated genes *TaNRT1, TaNRT2.1, TaNPF7.1*, and *TaGDH2* in transgenic wheat plants expressing either *OsPSTOL* (OsOE-3) or *TaPSTOL* (TaOE-1). Expression values shown are relative to the expression of *TaUbi* under in wheat plants grown under low nitrogen (low N) and replete nitrogen (replete) levels in hydroponic solution. Root specific expression is in the panels on the left and shoots specific expression in panels on the right for each gene. Data shown as the mean values (central line), lower and upper quartiles (box), minimum and maximum values (whiskers), and outliers as individual points. The statistical analysis was performed with ANOVA and post hoc Tukey test, letters correspond to significant differences between transcript levels of either line under either treatment (p⍰<⍰0.05).

## Discussion

To be able to translate our understanding of how plants tolerate various abiotic conditions it is important to understand if orthologous genes are present in various species or if the gene is part of a wider pathway. In this work we set out to understand why differences in the expression of *TaPSTOL* relative to *OsPSTOL* were seen (Gamuyao *et al*., 2012; Milner *et al*., 2018). To do this we created new transgenic material to make direct comparisons in a consistent genetic background and observe the transgene’s effects. From this we have learned that OsPSTOL can activate a conserved pathway in other cereal species and increase biomass production and ultimately yield under P limiting growth conditions (Figure 1). This suggests that wheat lacks a fully functional *PSTOL* type gene and that *TaPSTOL* is not the functional ortholog of OsPSTOL in wheat even though it is 92.7% similar (Milner *et al*., 2018). This is not that surprising as many different rice varieties including both reference varieties of japonica and indica also do not contain a functional *OsPSTOL* gene (Vigueira *et al*., 2016). The presence of a *PSTOL* type gene may have been selected in rice for under conditions for which wheat is not widely grown or has lost, as further mutations the near the N terminus of TaPSTOL locus reduced the N terminus of the protein. Or perhaps it was a chance event in rice and has never been present in wheat. This would fit with why no other direct ortholog of PSTOL has been found in any other species in the past decade (Bernardino *et al*., 2019; Milner *et al*., 2018).

It was found however that the benefits of overexpressing *OsPSTOL* in wheat conferred increased yields under a range of N conditions. This includes under N levels approximately 1/3 that of what current best practices in the UK for some transgenic lines (Figure 2). This mechanism of action seems to be by increasing biomass which leads to greater seed set. This was seen under both low P growth conditions as well as low N growth conditions (Figures 1 and 2). To further understand how *OsPSTOL* expression conferred tolerance to low N measurements of uptake, tissue concentrations and expression of N related genes were undertaken. From this it was found that *OsPSTOL* expression increased N uptake from the soil but did not change overall levels of N in the above ground tissues including the grain. This increased uptake of N was seen via differences in N regulated genes such as the high affinity uptake transporter *NRT2.1* the low affinity and N sensor *NRT1* and genes involved in N translocation *NPF7.1* (Figure 7) (Buchner and Hawkesford, 2014; Ho *et al*., 2009). This direct measurement of increased N uptake in plant overexpressing *OsPSTOL* is the first direct evidence that *OsPSTOL* increases uptake of nutrients to help aid growth.

This increased uptake did not change other physiological parameters including carbon assimilation or increased chlorophyll content suggesting that *OsPSTOL* helps plants take up or acquire more nutrients and does not alter perception of those nutrients unlike more recent reports involving altered brassinosteroid genes (Milner *et al*., 2021). We also show that *OsPSTOL* transcript levels can be seen under low N levels further supporting a role for *OsPSTOL* in nutrient uptake rather than just P directly. However this increase in transcript levels might be due to the cross talk of N and P in plants and not a direct activation of *OsPSTOL* by a N sensing transcription factor per se (Borah *et al*., 2018; Hong *et al*.,2012; Medici *et al*., 2019; Zhu *et al*., 2021).

Overall, it appears that *OsPSTOL* is able to aid in multiple nutrient deficiencies in different plant species by helping acquire the nutrients which are limiting growth. It is tempting to wonder if a combination of genes such as the recently cloned *OsNRT2.3b* or SPDT in combination with *OsPSTOL* could dramatically increase grain production under lower inputs (Fan *et al*., 2016; Yamaji *et al*., 2016).

## Data availability

Seed from transgenic lines is available upon request from the corresponding author. Seed materials will be transferred under MTA.

## Author contributions

MM and EW conceived the research. MM, MC and SB. performed experiments and MM wrote the manuscript and it was edited by EW.

## References

Azevedo, G.C., Cheavegatti-Gianotto, A., Negri, B.F., Hufnagel, B., e Silva, L. da C., Magalhaes, J. V, et al. (2015) Multiple interval QTL mapping and searching for PSTOL1 homologs associated with root morphology, biomass accumulation and phosphorus content in maize seedlings under low-P. BMC Plant Biol., 15, 172.

Bernardino, K.C., Pastina, M.M., Menezes, C.B., De Sousa, S.M., Maclel, L.S., Geraldo Carvalho, G.C., et al. (2019) The genetic architecture of phosphorus efficiency in sorghum involves pleiotropic QTL for root morphology and grain yield under low phosphorus availability in the soil. BMC Plant Biol., 19.

Borah, P., Das, A., Milner, M.J., Ali, A., Bentley, A.R., and Pandey, R. (2018) Long non-coding rnas as endogenous target mimics and exploration of their role in low nutrient stress tolerance in plants. Genes (Basel)., 9, 1.– 17.

Buchner, P. and Hawkesford, M.J. (2014) Complex phylogeny and gene expression patterns of members of the NITRATE TRANSPORTER 1/PEPTIDE TRANSPORTER family (NPF) in wheat. J. Exp. Bot., 65, 5697–5710.

Chin, J.H., Gamuyao, R., Dalid, C., Bustamam, M., Prasetiyono, J., Moeljopawiro, S., et al. (2011)Developing Rice with High Yield under Phosphorus Deficiency: Pup1 Seguence to Application. Plant Physiol., 156, 1202–1216.

Fan, X., Tang, Z., Tan, Y., Zhang, Y., Luo, B., Yang, M., et al. (2016) Overexpression of a pH-sensitive nitrate transporter in rice increases crop yields. Proc. Natl. Acad. Sci. U. S. A., 113, 7118–7123.

Gamuyao, R., Chin, J.H., Pariasca-Tanaka, J., Pesaresi, P., Catausan, S., Dalid, C., et al. (2012) The protein kinase Pstol1 from traditional rice confers tolerance of phosphorus deficiency. Nature, 488, 535–539.

Heuer, S., Lu, X., Chin, J.H., Tanaka, J.P., Kanamori, H., Matsumoto, T., et al. (2009) Comparative seguence analyses of the major quantitative trait locus p hosphorus up take 1 (Pup1) reveal a complex genetic structure. Plant Biotechnol. J., 7, 456–471.

Hiscox, J.D. and Israelstam, G.F. (1979) A method for the extraction of chlorophyll from leaf tissue without maceration. Can. J. Bot., 57, 1332–1334.

Ho, C.H., Lin, S.H., Hu, H.C., and Tsay, Y.F. (2009) CHL1 functions as a nitrate sensor in plants. Cell, 138, 1184–1194.

Hong, Y.F., Ho, T.H.D., Wu, C.F., Ho, S.L., Yeh, R.H., Lu, C.A., et al. (2012) Convergent Starvation Signals and Hormone Crosstalk in Regulating Nutrient Mobilization upon Germination in Cereals. Plant Cell, 24, 2857–2873.

Hufnagel, B., de Sousa, S.M., Assis, L., Guimaraes, C.T., Leiser, W., Azevedo, G.C., et al. (2014) Duplicate and Conquer: Multiple Homologs of PHOSPHORUS-STARVATION TOLERANCE1 Enhance Phosphorus Acquisition and Sorghum Performance on Low-Phosphorus Soils. Plant Physiol., 166, 659–677.

Magalhaes, J. V., Liu, J., Guimarães, C.T., Lana, U.G.P., Alves, V.M.C., Wang, Y.H., et al. (2007) A gene in the multidrug and toxic compound extrusion (MATE) family confers aluminum tolerance in sorghum. Nat. Genet., 39, 1156–1161.

Magnavaca, R., Gardner, C.O., and Clark, R.B. (1987) Evaluation of inbred maize lines for aluminum tolerance in nutrient solution. In: Genetic Aspects of Plant Mineral Nutrition, pp. 255–265. Dordrecht: Springer Netherlands.

Marschner, H. (1995) Mineral Nutrition of Higher Plants - 2nd Edition, 2nd edition. Academic Press.

Medici, A., Szponarski, W., Dangeville, P., Safi, A., Dissanayake, I.M., Saenchai, C., et al. (2019) Identification of Molecular Integrators Shows that Nitrogen Actively Controls the Phosphate Starvation Response in Plants. Plant Cell, 31, 1171–1184.

Milner, M.J., Howells, R.M., Craze, M., Bowden, S., Graham, N., and Wallington, E.J. (2018) A PSTOL-like gene, TaPSTOL, controls a number of agronomically important traits in wheat. BMC Plant Biol., 18, 115.

Milner, M.J., Swarbreck, S.M., Craze, M., Bowden, S., Griffiths, H., Bentley, A.R., and Wallington, E.J. (2022) Over-expression of TaDWF4 increases wheat productivity under low and sufficient nitrogen through enhanced carbon assimilation. Commun. Biol. 2022 51, 5, 1–12.

Milner, M.J., Swarbreck, S.M., Craze, M., Bowden, S., Griffiths, H., Bentley, A.R., and Wallington, E.J. (2021) Over-expression of the brassinosteroid gene TaDWF4 increases wheat productivity under low and sufficient nitrogen through enhanced carbon assimilation. bioRxiv, 2021.09.21.461226.

Vigueira, C.C., Small, L.L., and Olsen, K.M. (2016) Long-term balancing selection at the Phosphorus Starvation Tolerance 1 (PSTOL1) locus in wild, domesticated and weedy rice (Oryza). BMC Plant Biol., 16, 101.

Wissuwa, M., Wegner, J., Ae, N., and Yano, M. (2002) Substitution mapping of Pup1⍰: a major QTL increasing phosphorus uptake of rice from a phosphorus-deficient soil. TAG Theor. Appl. Genet., 105, 890–897.

Xu, K., Xu, X., Fukao, T., Canlas, P., Maghirang-Rodriguez, R., Heuer, S., et al. (2006) Sub1A is an ethylene-response-factor-like gene that confers submergence tolerance to rice. Nature, 442, 705–708.

Yamaji, N., Takemoto, Y., Miyaji, T., Mitani-Ueno, N., Yoshida, K.T., and Ma, J.F. (2016) Reducing phosphorus accumulation in rice grains with an impaired transporter in the node. Nat. 2016 5417635, 541, 92–95.

Zhu, Z., Li, D., Cong, L., and Lu, X. (2021) Identification of microRNAs involved in crosstalk between nitrogen, phosphorus and potassium under multiple nutrient deficiency in sorghum. Crop J., 9, 465–475.

